# A translational transcriptomic signature of vaccine reactogenicity for the evaluation of novel formulations

**DOI:** 10.1101/2025.11.20.689650

**Authors:** Jérémie Becker, Maroussia Roelens, Kendra Reynaud, Laurent Beloeil

**Author notes:** Correspondence & lead contact: Further information and requests for resources should be directed to and will be fulfilled by the corresponding author, Laurent Beloeil. These authors contributed equally to this work. These senior authors contributed equally to this work. See group listing in the collaborators section.

## Abstract

Accurately predicting vaccine reactogenicity at the preclinical stage remains a major challenge in vaccine development, as conventional animal studies and in vitro assays capture general inflammation but fail to quantify local or systemic reactogenicity relevant to humans. Using transcriptomic data from the BioVacSafe consortium encompassing seven vaccines and immunostimulants in mice and five licensed vaccines in humans, we developed a cross-compartment and cross-species predictive model of vaccine reactogenicity. Reactogenicity classes were defined in mouse muscle based on the magnitude of transcriptomic responses and literature evidence. A penalized ordinal regression model was trained to predict both discrete classes and continuous scores of reactogenicity. Transcriptomic profiles from mouse muscle were highly predictive of reactogenicity, and the most predictive genes pointed to inflammatory and tissue repair programs such as IL6/JAK/STAT3 signalling. When transferred to mouse blood, the model reliably separated high-from low-reactogenicity formulations, although the intermediate class was less well resolved, suggesting shared transcriptional programs between compartments and coordinated innate responses. When applied to human blood, the classifier identified Fluad (MF59-adjuvanted) as the most reactogenic licensed vaccine, in agreement with elevated C-reactive protein and ReactoScore values, whereas smaller differences among the remaining vaccines were not resolved. These results align with clinical safety data and suggest that early transcriptomic signatures in mice can identify highly reactogenic formulations in humans. Our study presents a pan-vaccine, cross-species transcriptomic signature that bridges preclinical and clinical data, offering a foundation for translational biomarkers and mechanism-informed screening of vaccine tolerability.

## INTRODUCTION

Vaccine reactogenicity refers to the local and systemic clinical manifestations that occur after vaccination, partly driven by inflammation resulting from the recruitment and activation of immune cells. Although most events are transient and mild, their frequency and severity vary across vaccine platforms (e.g., adjuvanted protein, viral vector, mRNA) and between individuals^1^. This variability complicates both clinical assessment and regulatory decision-making^2^.

The COVID-19 pandemic highlighted the need for reliable approaches to evaluate vaccine reactogenicity not only prior to clinical testing but also during vaccination campaigns, where responses have been shown to vary according to age, sex, and pre-existing comorbidities^3^. Across the vaccine development pipeline, reactogenicity and safety are monitored, beginning with *in vitro* assays and animal toxicology studies^4^, progressing through clinical trials, and extending to post-licensure pharmacovigilance^2^. Preclinical evaluations are intended to exclude highly reactogenic candidates early, thereby reducing costs and focusing resources on the most promising formulations. Nevertheless, the mechanisms underlying local and systemic symptoms after vaccination remain poorly understood^3^, and no reliable biomarkers have yet been established to predict human reactogenicity^2^. The discovery of translatable markers of reactogenicity therefore remains an active area of research.

To address these limitations, translational research increasingly relies on preclinical models and microphysiological platforms, including organs-on-chips and *in vitro* assays^5^, to elucidate the mechanisms of action of vaccines and evaluate their associated adverse events. For example, the Modular Immune In vitro Construct platform^6^ predicts local and systemic adverse events by profiling cytokine and chemokine release from PBMCs. Model performance is driven by four mediators (IL-1β, IL-6, IL-10, and CCL4), and when applied to Trumenba, a vaccine not included in training, the model accurately ranked observed local and systemic reactogenicity. While representing the first quantitative framework for reactogenicity prediction, the current results remain constrained by the lack of paired biological and clinical data as well as its reliance on a limited panel of markers.

Preclinical studies in mice and rabbits, on the other hand, provide information on innate immune activation, cytokine release, and physiological endpoints, with rabbits serving as the regulatory standard for vaccine toxicity assessment^4^. These studies typically rely on markers such as IL-6, CRP, hepatic enzymes, and histopathology^5^. While informative, such endpoints mainly capture general inflammatory responses rather than specific features of vaccine reactogenicity. In comparative adjuvant studies in adults receiving hepatitis B antigen formulated with AS01B or Alum, elevated IL-6 and IFN-γ levels were associated with systemic reactogenicity following AS01B administration^7^. Similarly, findings from the BioVacSafe studies revealed conserved interferon and chemokine-related transcriptional modules such as IL-10 and CXCL10, that were induced in both mouse and human datasets^8,9^. Yet, these cross-species commonalities remain largely descriptive and do not provide quantitative information on the magnitude or type (local versus systemic) of reactogenicity that may occur in humans. Overall, these studies underscore both the value and the limitations of early-phase biomarkers and highlight the need for improved translational models that bridge preclinical readouts with clinical tolerability outcomes.

The present study builds on BioVacSafe, an Innovative Medicines Initiative-funded public-private partnership aimed at systematically identifying biomarkers of relatively common inflammatory events after adjuvanted immunization using human, animal, and population-based models^10^. This consortium pioneered harmonized multi-tissue transcriptomics and correlation-based analyses to identify biomarkers of vaccine reactogenicity across species^8,9^. Although these studies provided key mechanistic insights into vaccine reactogenicity, to our knowledge no predictive signature (*i.e.* set of genes) has yet been reported, either within the consortium or by other groups. To address this gap, we adopt a predictive framework aimed at identifying translational biomarkers of reactogenicity from animal to human. Based on BioVacSafe transcriptomic data (**Figure 1a,b**), we first derive a time-specific signature of reactogenicity from mouse muscle following exposure to four licensed vaccines with established safety profiles and three immunostimulants with defined inflammatory potentials (**Figure 1d**). The diversity of these formulations provides a robust basis to capture genuine reactogenicity patterns and supports generalization to a broad range of vaccine candidates. We then tested this reactogenicity signature in mouse blood to assess its applicability in a more accessible compartment (**Figure 1d**), facilitating future biomarker studies. Finally, we applied the reactogenicity signature to human blood across additional vaccine formulations to evaluate cross-species translatability. This three-stage strategy assesses whether signatures identified in mouse muscle can be detected in mouse and human blood, and thus whether animal data can inform vaccine safety assessment in clinical settings. Of note, to train the signature in mouse muscle, reactogenicity classes are defined by combining the magnitude of the transcriptomic response in mice with evidence from the human literature. For simplicity, we therefore retain the term “reactogenicity” for both species, even though mouse models are not typically used to define it. Furthermore, the terms signature, classifier and model are used interchangeably in the rest of the paper.

**Figure 1:**
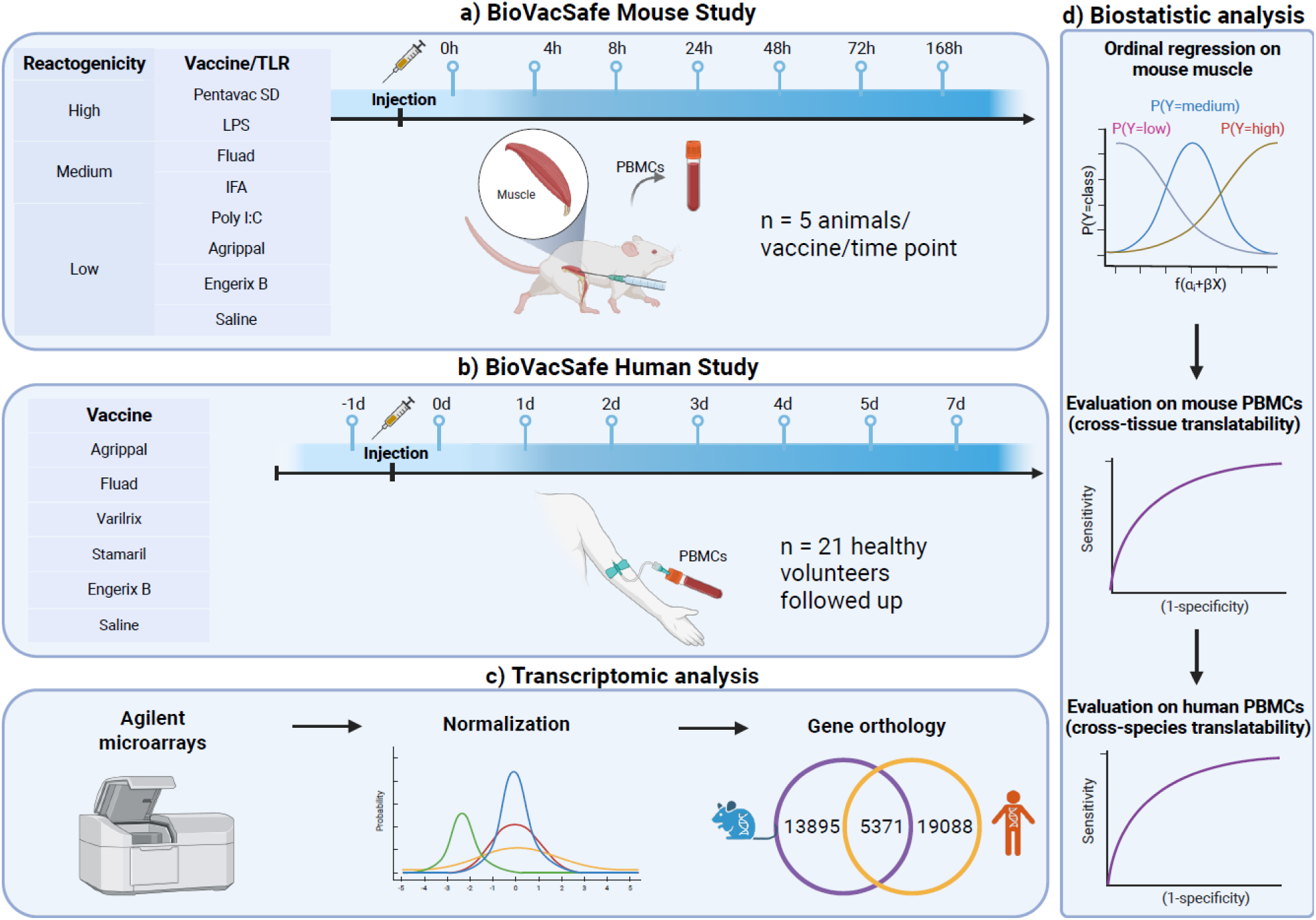
**Study analysis workflow**. BioVacSafe **(a)** mouse and **(b)** human transcriptomic datasets were retrieved, **(c)** normalized, and mapped through orthologous probes to enable cross-species comparison. A penalized ordinal regression model was trained on mouse muscle data to derive a time-specific reactogenicity signature. **(d)** The signature was subsequently applied to mouse and human blood to assess its translatability across compartments and species.

## RESULTS

### Defining Reactogenicity Classes from Transcriptomic and Literature Evidence

In the absence of head-to-head clinical reactogenicity data for the seven formulations, no foundational dataset was available to train the predictive model. We thus established a provisional hierarchy by using the first principal component of mouse muscle as proxy for inflammation, assuming that inflammatory response represented the main source of variation during the early post-vaccination period. This inferred hierarchy was then reconciled with published vaccine literature to define reactogenicity classes. These classes were subsequently used to train a penalized ordinal regression model on mouse muscle, the compartment in which vaccine response was found to be strongest in the original study^8^. The fitted model was then applied on the three transcriptomic datasets (mouse muscle and blood, human blood), to predict a continuous reactogenicity score, hereafter referred to as the *latent variable* (see **Materials and Methods**), as well as reactogenicity classes obtained by discretizing this latent variable.

In this way, the effects of time and vaccine formulation were examined along the first two principal components (PCs) in each dataset (**Supplementary Figure 1**). Although the proportion of explained variance captured by these PCs was low in all four datasets, distinct patterns emerged. In mouse muscle, vaccine formulations were broadly distributed along the first principal axis in the following order: Saline, Agrippal, Engerix B, Poly I:C, Fluad, IFA, LPS and Pentavac SD. This gradient suggested that the formulations elicited qualitatively similar responses that differed mainly in magnitude. The second axis captured temporal variation, with values increasing across successive time points. Thus, both biological differences between formulations and temporal dynamics were detected, despite the relatively low explained variance. A comparable, though less pronounced, pattern was observed in mouse blood: LPS and Pentavac SD remained distinct from other vaccines; early (4–24h) and late (48–168h) time points exhibited differing directions. In the combined mouse dataset, the two compartments (muscle versus blood) were separated along a direction that was mostly aligned with the first principal component. As all values are expressed relative to the pre-vaccination baseline of the same vaccine group within the same compartment, this separation did not reflect differences in baseline expression between muscle and blood, but rather differences in the amplitude and kinetics of the response between compartments. Vaccine formulations were distributed along a trajectory following approximately the second component, and the ellipses showed the same orientation in both compartments, indicating that the vaccine effect was conserved across tissues. In contrast, the human dataset showed substantial overlap across both vaccine types and time points, pointing to a weaker signal.

When examined at individual time points, the two mouse datasets recapitulated a similar hierarchy with clearer separations than in the joint analysis (**Supplementary Figure 2**). As already noted in the initial study^8^, this hierarchy was not strictly conserved over time due to distinct kinetics among formulations. Because our goal was to capture the overall hierarchy of reactogenicity rather than time point-specific differences, we derived a classification that remained broadly stable and homogeneous across time points. Owing to the limited availability of direct comparisons between formulations in the literature, we grouped them along the first principal component into three classes: low (Poly I:C, Agrippal, Engerix B and Saline); medium (IFA and Fluad), and high (LPS and Pentavac SD). The high- and low-reactogenicity classes were separated at all time points up to 72h in both mouse muscle and mouse blood, while the three classes were most distinct at 24 and 48h in mouse muscle (**Supplementary Figure 2**). In mouse blood, the medium class overlapped with the low class, in line with the observations on aggregated time points (**Supplementary Figure 1**).

Although reactogenicity levels of vaccine formulations varies considerably between humans and mice due to species-specific immune mechanisms and differences in vaccine/adjuvant properties^11^, those three classes observed in mouse muscle are consistent with reactogenicity profiles described in the literature. LPS, a canonical inducer of systemic inflammation in mice, is among the most reactogenic agents^12^. In humans, LPS toxicity has precluded direct use, leading to detoxified derivatives such as monophosphoryl lipid A, which retains immune stimulation while reducing harmful effects^13^. Reactogenicity in Pentavac SD is driven by its whole-cell pertussis component, which is inherently more reactogenic than acellular alternatives^14^.

Among the medium-reactogenicity formulations, Montanide (IFA-type) emulsions have been demonstrated to reliably boost immunogenicity and are characterized by dose- and formulation-dependent local reactogenicity such as commonly pain, nodules or granulomas^15–17^. In clinical trials using Montanide with peptide antigens (*e.g.* HIV and malaria vaccines), sterile abscesses were reported, limiting their broader prophylactic use^18^. Compared with non-adjuvanted influenza vaccines, MF59-adjuvanted formulations were shown to have higher rates of mild-moderate reactions at local injection site but comparable AEs^19^.

Within the low-reactogenicity class, Poly I:C adjuvantation was investigated in human vaccine trials and glioblastoma settings, where it was described as safe and well tolerated, with predominantly low-grade local reactions (*e.g.* mild injection site effects) and no clear pattern of severe systemic toxicity^20,21^. Across 36 clinical trials, Engerix-B displayed modest adverse event rates, with injection-site soreness reported in ∼22% of recipients and fatigue in ∼14%, confirming its overall favourable tolerability profile^22^.

### Transcriptomic profiles in mouse muscle are highly predictive of vaccine reactogenicity

Building upon these reactogenicity classes, we trained and evaluated an ordinal regression model on mouse muscle. These two steps were applied to both the complete mouse microarray and the subset of probes corresponding to genes orthologous to human genes. The weighted F1 score, combining precision and recall across classes, indicated that all six time points were predictive of reactogenicity class, with peak performance observed at 8h and 24h in the full set (median weighted F1 = 0.98; **Figure 2a**). This was consistent with the greater class separation seen at 24h in the PCA (**Supplementary Figure 2**). Interestingly, the median F1 scores were identical between the two probe sets across all time points except 8h, suggesting that the reactogenicity signal was largely preserved in orthologous genes. To contextualize these results in terms of classification error, confusion matrices were generated. Across 100 cross-validation repetitions, the average number of misclassified samples, out of 40 samples per time point, ranged from one at 24h to more than three at 4h and 168h (**Supplementary Figure 3a**), corroborating the higher predictive performance at 24h.

**Figure 2:**
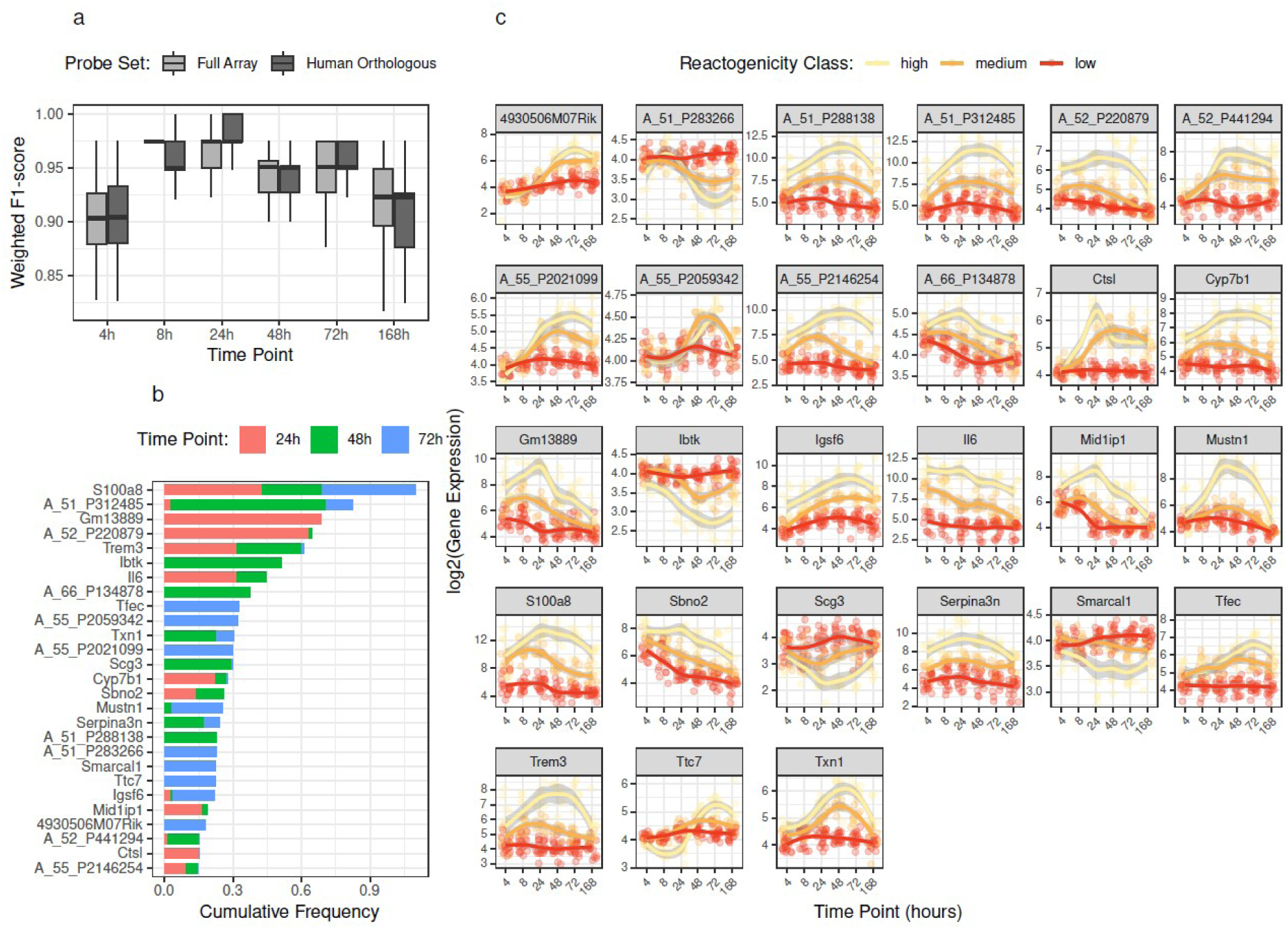
Pan-vaccine signature definition and evaluation in mouse muscle. **(a)** A penalized ordinal regression model was trained and evaluated using repeated nested cross-validation at each time point, with performance assessed by the weighted F1-score on both the full mouse microarray and the subset of probes orthologous to human genes. **(b)** Gene importance for predicting reactogenicity was assessed by stability selection. The top 10 most frequently selected genes at 24, 48, and 72 hours are displayed. **(c)** Expression trajectories of the top 27 genes are shown over time for each reactogenicity class.

We next sought to characterize the main drivers of the reactogenicity signature, i.e., genes with large selection frequency (**Supplementary Table 1**). We first conducted a differential analysis between high and low classes to quantify effect size and assess statistical significance (**Supplementary Table 2**). Although log_2_ fold-change did not correlate globally with selection frequency, reflecting the distinct statistical properties of univariate and multivariate analyses, the comparison revealed that the most predictive genes (frequency > 10%) almost systematically exhibited |log_2_ fold-change| > 1 and adjusted p-values < 10^-5^, thresholds commonly considered biologically meaningful (**Supplementary Figure 3b**). Visual inspection of the top 10 most frequently selected genes at each time point (**Figure 2b**) confirmed that these 27 unique genes underwent substantial modulation in one or more reactogenicity classes, with log₂ fold-changes reaching up to 2 (**Figure 2c**). The relative ordering of classes was preserved for most genes, with some exceptions (e.g., *Ctsl*), indicating that the ordinal regression model largely identified genes associated with graded vaccine reactogenicity. Most genes showed maximal class separation between 24 and 72 hours, consistent with the classification performance trends described above.

Additionally, we performed a pathway enrichment analysis on the selection frequency at each time point. Although none of MSigDB Hallmark gene sets reached statistical significance (p-values > 0.12), the seven pathways highlighted by GSEA aligned well with inflammatory processes such as TNFα Signalling via NF*-* κB, Inflammatory Response and IL-6/JAK/STAT3 signalling (**Supplementary Figure 3c)**. Furthermore, IL-6, which was among the top 10 most selected gene at 24h, has been reported to be a key component of vaccine-induced reactogenicity in the literature^6,7,23^. Epithelial–Mesenchymal Transition was also highlighted by pathway enrichment, and is known to play a role in tissue repair^24^, suggesting that reactogenicity prediction does not solely rely on inflammatory markers, reflecting the general significance of inflammation and of acute phase reactants^25^.

### The reactogenicity signature translates across compartments

Following the identification of a pan-vaccine classifier in mouse muscle, we applied the model to mouse blood to evaluate its translatability across compartments. This step was motivated by the predominance of blood-based immune monitoring in vaccine studies and by the practical consideration that any clinically deployable signature would need to be measurable in peripheral blood, where sampling is far less invasive than muscle biopsies.

The median weighted F1 ranged from 0.57 to 0.80, *i.e.* up to 0.35 lower than the corresponding performance in muscle at 24h and 48h (**Supplementary Figure 4a**). Despite this decrease, examination of Figure 3a revealed that Pentavac SD and LPS consistently exhibited higher median latent variables (*i.e.* continuous reactogenicity scores) than the other six vaccines across time-point combinations, with the most pronounced differences observed when applying the model at 72h in blood (see pairwise t-test in **Supplementary Figure 4d**). By contrast, the medium class showed minimal separation from the low-reactogenicity formulations, a finding consistent with PCA trends. These results indicate that the transition from muscle to blood primarily impacts the distinction between low and medium reactogenicity groups, a pattern also visible in the dynamics of gene expression (**Figure 3b**). Given that the intermediate reactogenicity class likely reduces model performance, we retrained the model in muscle using low and high reactogenicity classes and reapplied it to blood data. This refined model achieved better separation, with a median weighted F1 = 0.95 across all time points (**Supplementary Figure 4a**), pointing to robust translatability of the model to mouse blood when the intermediate class was excluded. Of note, classification was maximal (weighted F1 = 1.00) at 72 h in muscle (**Supplementary Figure 4a**), the time point at which the log_2_ fold-change between low and high reactogenicity across compartments correlated most strongly (R = 0.59, **Supplementary Figure 4b**). These higher classification and correlation values found at 72 h suggest that, despite differences in the magnitude and kinetics of transcriptomic profiles across muscle, blood, and lymph nodes^8^, the responses in muscle and blood exhibit the greatest similarity at this time point. These observations point toward shared transcriptional programs between compartments, potentially driven by immune cells trafficking from the injection site to the circulation.

**Figure 3:**
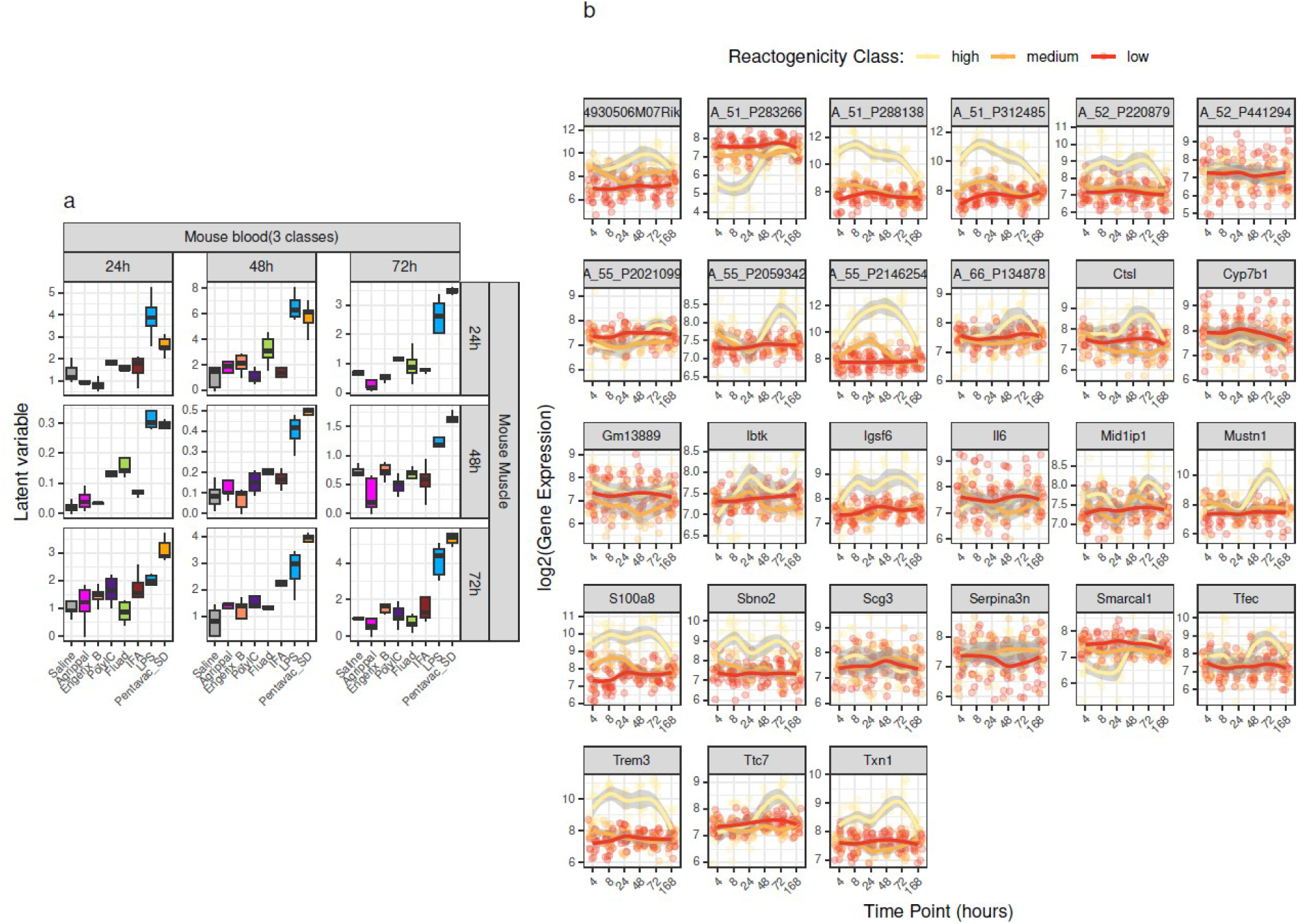
Predicted reactogenicity in mouse blood. **(a)** Latent variables predicted across all combinations of three time points (24h, 48h, 72h), with models trained in mouse muscle and applied in mouse blood. **(b)** Temporal expression profiles of the most predictive genes for each reactogenicity class in mouse blood.

### The reactogenicity signature translates across species and is consistent with inflammation readouts

We finally investigated whether the pan-vaccine classifier generalize across species by applying it to human blood using both two-class and three-class models. In the absence of comparative clinical studies directly assessing vaccine reactogenicity side by side, as noted earlier, and given that PCA revealed no clear separation between formulations in human blood, the definition of reactogenicity classes was consequently not performed in humans. Instead, we assessed the consistency between our model predictions and inflammation readouts (ReactoScore and CRP). Despite smaller variability across formulations than in mouse blood, Fluad displayed higher values than the other vaccines in CRP, ReactoScore, and latent variables predicted from mouse muscle at 24h (**Figure 4**). These differences were significant for both inflammation readouts and for the two-class model applied to human blood at 24h (**Supplementary Figure 5**). By contrast, only CRP and ReactoScore exhibited higher levels for Agrippal and Engerix-B than for Saline, Varilrix, and Stamaril across all time points, a pattern not captured by the model. Despite the absence of gold standard, the increased predicted reactogenicity in Fluad is consistent with reported safety profiles. As mentioned earlier, in a meta-analysis of 24 studies, Domnich *et al.* concluded that MF59-adjuvanted flu vaccines are generally more immunogenic, induce more reactogenic events, without increasing serious adverse events^19^. Stamaril^26,27^, Varilrix and Engerix B^29^ on the other hand are widely regarded as having good safety profiles. Taken together, these findings show that the model identified the most reactogenic licensed vaccine, in agreement with inflammation readouts and clinical safety data, whereas the non-significant trend towards higher CRP and ReactoScore values for Agrippal and Engerix-B was not captured. This supports its potential as a preclinical tool for flagging formulations with a high inflammatory profile.

**Figure 4:**
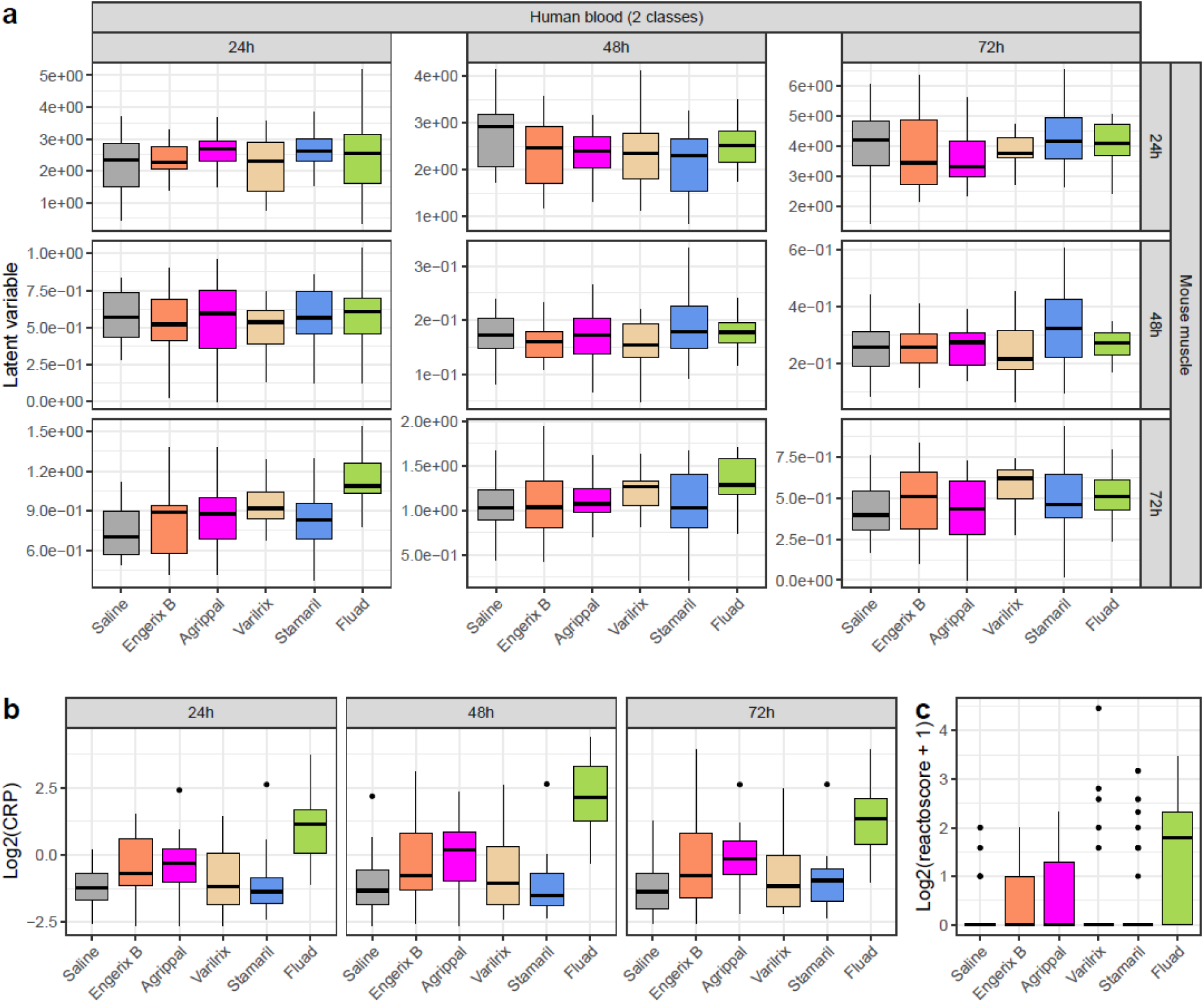
Predicted reactogenicity in human blood. **(a)** Latent variables predicted across all combinations of three time points (24h, 48h, 72h), using models trained in mouse muscle and applied in human blood. **(b)** CRP and **(c)** ReactoScore measured in the same individuals for comparison.

## DISCUSSION

Despite their central role in vaccine safety, the cellular and molecular mechanisms of vaccine reactogenicity remain incompletely understood. Upon vaccination, clinical signs related to reactogenicity are observed both locally and systemically. Clinical symptoms are hard to predict prior to injection, because they are a product of various direct and indirect mechanisms that elicit different reactions^2^. A major outlying question in the field is how diverse molecular mediators that are detected at different time points post vaccination translate into clinical events.

Preclinical assessment of vaccine reactogenicity is therefore critical to ensure safety before first-in-human trials. Yet current practice relies on a narrow set of surrogate markers, such as hepatic enzymes for toxicity or CRP in rabbits for inflammation^30^. The rapid deployment of mRNA vaccines during the COVID-19 pandemic further exposed the limitations of these tools, underscoring the need for predictive, mechanism-informed biomarkers that can be integrated early in vaccine development^10,31^.

To overcome this limitation, we developed a pan-vaccine transcriptomic signature of reactogenicity by leveraging data from the BioVacSafe consortium. We first focused on the injection site, where transcriptional changes are stronger than in peripheral blood^8^ and where diverse immune cells are rapidly recruited upon vaccination^32–34^. Licensed vaccines and immunostimulants were classified into three reactogenicity categories based on PCA results and prior knowledge of their inflammatory profiles. From mouse muscle, we identified a predictive signature that remained robust when transferred to blood after excluding the intermediate reactogenicity class. This transfer, strongest at 72h where muscle and blood responses were most concordant, is consistent with a local inflammatory response at the injection site giving rise to a detectable systemic signal. When applied to human data, the classifier correctly identified Fluad as the most reactogenic licensed vaccine, consistent with CRP levels, ReactoScore, and established clinical safety profiles. The wide diversity of formulations used in mice, from saline to LPS, provides a unique advantage: the model was trained on across a reactogenicity spectrum wider than that observed among licensed human vaccines, enabling it to predict whether a new formulation lies within a “tolerability zone” or exceeds it. Collectively, these findings demonstrate the potential of a cross-compartment, cross-species framework, agnostic to vaccine platform.

Although marked differences exist between mice and humans in pattern-recognition receptor expression, signalling pathways, immune cell subset composition, cytokine responses and regulatory networks^35^, the transfer of the signature to mouse blood and its identification of the most reactogenic vaccine in human blood, together with the diversity of formulations in the training set, suggest that the model captures host-response mechanisms conserved across species. These results also support the notion that key components of vaccine-induced inflammation and reactogenicity are evolutionarily conserved. Viewing reactogenicity as part of a homeostatic process^36^ may explain why predictive patterns of innate activation are preserved across species^37,38^. In summary, when applied to licensed vaccines with well-established safety profiles in human, our signature identifies the licensed vaccine with the highest CRP response and can be implemented in mouse models to infer vaccine safety at the preclinical stage. Importantly, unlike CRP which increases only moderately in mice compared to humans^39^ and whose complement-activating role is not conserved across species^39,40^, our transcriptomic signature turned out to be translatable.

Similar studies have reported predictive transcriptomic signatures of vaccine outcomes. For immunogenicity, both baseline^41^ and post-vaccination^42^ markers have been shown to effectively predict antibody responses, the former being derived from thirteen vaccines. For reactogenicity, an infant study of 4CMenB identified four post-vaccination markers of febrile responses in plasma proteins and was subsequently validated in mice^43^. Our work extends the latter by providing a pan-vaccine, cross-species signature of reactogenicity based on early post-vaccination responses, demonstrating translational potential in preclinical safety evaluation.

Several limitations should be noted. First, the model was deliberately trained on a reactogenicity range wider than that of licensed vaccines, from saline to LPS. This favours the detection of strongly reactogenic formulations but limits resolution among vaccines with moderate profiles: in human blood, the model identified Fluad but did not recover the trend towards higher CRP and ReactoScore values for Agrippal and Engerix-B compared with the other non-Fluad vaccines. Second, reactogenicity classes were derived from the first principal component of mouse muscle, which explains a modest share of variance, and were reconciled with literature evidence in the absence of head-to-head clinical data; sensitivity analyses across alternative normalization and dimensionality reduction strategies were not performed. Third, the signature was derived from muscle as a proxy for local reactogenicity, and its relevance to systemic reactogenicity rests on the transcriptional programs shared with blood. The human comparison provides only limited evidence on this point, as the retained ReactoScore is dominated by injection-site reactions and CRP is the only purely systemic readout available.

As a next step, we envision validating the current signature across a broader panel of vaccine formulations and in human cell-based models to reinforce its functional relevance and predictive value^44^. An additional line of development will involve assessing its applicability across emerging vaccine platforms, including viral vectors, mRNA formulations, and novel adjuvants such as polyacrylate^45^, thereby evaluating its robustness in next-generation vaccine development. A complementary analysis will consist of retraining the model without the most extreme stimuli (LPS and Pentavac SD) to assess whether finer gradations among moderately reactogenic vaccines can be recovered. In parallel, a discovery analysis performed directly in blood could identify reactogenicity-associated genes that are not part of the muscle-derived signature and that may be more directly deployable for clinical monitoring. A subsequent signature reduction step, retaining only the most informative transcripts, will further enable its translation to low-throughput platforms such as multiplex qPCR. Ultimately, the signature could also be refined for specific populations, such as paediatric or immunocompromised individuals, enabling more accurate and clinically relevant assessment of vaccine reactogenicity.

## MATERIAL & METHODS

### BioVacSafe datasets

As described in the original study^8^, seven vaccines and immunostimulants were evaluated in the mouse study : Pentavac SD (Diphtheria, tetanus, pertussis (whole cell), hepatitis B (rDNA) and haemophilus type b conjugate vaccine), Agrippal (Trivalent flu subunits – H3N2, H1N1 and influenza B), Fluad (Trivalent flu subunits – H3N2, H1N1 and influenza B + MF59), Engerix B (Recombinant hepatitis B sAg absorbed on alum), IFA (Montanide ISA 51 VG), LPS (LPS-EB Ultrapure), and the TLR3 agonist Poly I:C (Polyinosinic:polycytidylic acid), alongside a saline control. Mice were culled at six time points post-injection (4h, 8h, 24h, 48h, 72h, and 168h) and an unvaccinated (0h) group was included for each treatment. The injected muscle site and lymph nodes draining the hind leg quadriceps muscles were harvested. Peripheral blood was sampled from the mouse tail vein. Total RNA was isolated, labelled, hybridized and profiled on an Agilent microarray. Each vaccine group and time point included five animals.

The human study^9^ assessed five licensed vaccines, Agrippal, Fluad, Engerix B, Stamaril (live-attenuated YFV vaccine), Varilrix (live-attenuated varicella zoster virus) along with a saline control, the first three overlapping with those tested in mice. Participants were followed longitudinally, peripheral blood was collected from one day before vaccination through 28 days post-vaccination. Only samples obtained up to seven days post-vaccination (D-1, D0, D1, D2, D3, D4, D5, D7) were included to match the time points in the mouse study. Transcriptomic profiling was performed on blood-derived RNA and inflammatory mediators, including C-reactive protein (CRP), were quantified. Adverse events (AEs) were systematically documented, including description (e.g. headache, pain at injection site), classification (e.g. nervous system disorders, general disorders and administration site conditions), severity grading (mild, moderate, or severe), and duration (in days). AEs deemed non-specific (e.g. fatigue, nausea, nasal congestion, sleep disorders etc), unrelated to vaccination, recorded after 7 days post-vaccination, associated with indwelling cannula, or arising from laboratory artifacts were excluded from the analysis. The list of AEs retained in the analysis is available **Supplementary Table 3.** Similar to the original study^9^, a ReactoScore was calculated for each patient as follows:

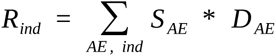

where *S_AE_* is the AE severity (1 = mild, 2 = moderate, 3 = severe), and *D_AE_* its duration in days. Of note, patients with no documented AE were assigned a ReactoScore of 0.

### Data preprocessing

Processed expression data were retrieved from the NCBI Gene Expression Omnibus (GEO) portal ^44,45^ using the R package *GEOquery* (accession numbers GSE120661 and GSE124533), as background-corrected and normalized by the original BioVacSafe investigators with the R package *limma*. Background correction was performed with the normexp method for the mouse arrays^8^, whereas the human arrays relied on the background-corrected signal (gProcessedSignal) from Agilent Feature Extraction^9^. In both datasets, arrays were then quantile normalized. Starting from these log2 normalized intensities, control probes were removed, as well as probes lacking any gene annotation (gene symbol or Ensembl ID for the mouse array; gene symbol, Entrez Gene ID or Ensembl ID for the human array). Only probes corresponding to protein-coding transcripts, as annotated in Ensembl Biomart, were retained (matched on Ensembl transcript ID for the mouse array, and on gene symbol then Ensembl transcript ID for the human array). Replicate probes were averaged (*avereps*). Lowly expressed probes were then filtered within each dataset, retaining probes whose summed expression across samples exceeded a dataset-specific threshold multiplied by the number of samples in the smallest treatment–time point group. For the human dataset, only samples collected up to day 7 were retained, and the Engerix B three-dose arm and its corresponding placebo arm were excluded.

For each dataset (mouse muscle, mouse blood, human blood) and each vaccine group, a baseline profile was computed as the gene-wise median of the pre-vaccination samples of that group (0h in mice, D-1 in humans) and subtracted from every sample of the group, on the log2 normalized intensity scale. All downstream analyses, except the differential analysis, were performed on these baseline-corrected values, which therefore represent log2 changes relative to the pre-vaccination state of the same vaccine group within the same compartment. A group-median baseline was used for the human data as well, rather than a within-subject reference, to apply a uniform correction across datasets. Because mice were culled at each time point, the mouse baseline is necessarily a group-level reference.

### Orthology between mouse and human Agilent microarrays

GPL annotation files for the mouse and human Agilent microarrays were retrieved from GEO. Probe coordinates were based on the MGSCv37^48^ (mouse) and GRCh37^49^ (human) genome assemblies. Mouse probe positions were converted to GRCh37 using UCSC LiftOver^50^. Both the converted mouse probes and the original human probes were then mapped to exons from the GRCh37 GTF. A mouse–human probe pair was defined as orthologous if both mapped to at least one common exon. After excluding one-to-many matches, 5,371 orthologous probe pairs were retained.

### Principal component analysis

Principal Component Analysis (PCA) was performed separately on each dataset, as well as jointly on the mouse muscle and blood data, to enable comparison of vaccine effects across compartments. To make the comparison possible, the microarrays were further quantile normalized together. For each dataset, sample projections were coloured by vaccine or time points and ellipses associated with reactogenicity classes were added.

### Differential analysis

Transcriptomic data from mouse muscle were analyzed for differential expression using the *lmFit* function of the R package *limma*. As in the original study, models including all interactions between treatments and time points were fitted. Instead of constructing contrasts comparing each vaccine against pre-vaccination and saline controls at each time point, comparisons were made at the class level. Specifically, we contrasted the high-reactogenicity class (LPS and Pentavac SD) with the low-reactogenicity class (Engerix B, Agrippal, Poly I:C, and Saline) using double contrasts (H _t_ – H_0h_) – (L_t_ – L_0h_), where *H* denotes the high-reactogenicity group, *L* the low-reactogenicity group, and *t* the considered time point. P-values were adjusted for multiple testing using Benjamini– Hochberg correction^51^.

### Predictive model

To predict reactogenicity from gene expression data, a penalized ordinal regression (*glmnetcr* R package) was applied to account for the ordered nature of the reactogenicity classes (**Figure 1d**). The model incorporates an L1 penalty to enforce sparsity and enable variable selection. A key feature of ordinal regression is that it models the response variable as a discretized version of an unobserved continuous latent variable which we use as a continuous proxy for reactogenicity. The cut-points defining the class boundaries are ordered in a strictly increasing manner and are estimated along the regression coefficients. The model was first trained and evaluated on mouse muscle data, where coefficients and penalization level were estimated using 5-folds nested cross-validation. The cross-validation procedure was repeated 100 times to obtain a stable estimate of model performance and to quantify uncertainty in the performance metrics. To address class imbalance, observation weights were applied so that each class contributed equally to parameter estimation. The *Caret* R package was further used to generate balanced folds. The estimated coefficients were then applied on mouse and human blood data to compute latent variables and reactogenicity predictions. Model performance was evaluated using the weighted F1 score, which balances precision and recall across classes while accounting for their relative frequencies, providing a robust measure in the presence of class imbalance. Weighted F1 score ranges from 0 to 1, where values close to 0 indicate poor classification performance while values approaching 1 indicate high precision and recall across all classes, reflecting excellent predictive performance. To identify the support of the molecular signature, variable importance was measured using stability selection^52^, a method based on subsampling.

To functionally characterize the signature, we performed a pathway enrichment analysis on gene selection frequencies using GSEA^53^, assessing the association between gene ranking and MSigDB Hallmark gene sets.

## Code availability

## ACKNOWLEDGMENTS

We gratefully acknowledge Prof. Giuseppe Del Giudice and Prof. Alberto Mantovani for their input regarding the BioVacSafe datasets and helpful discussions on vaccine reactogenicity.

---

**Supplementary Figure 1:**
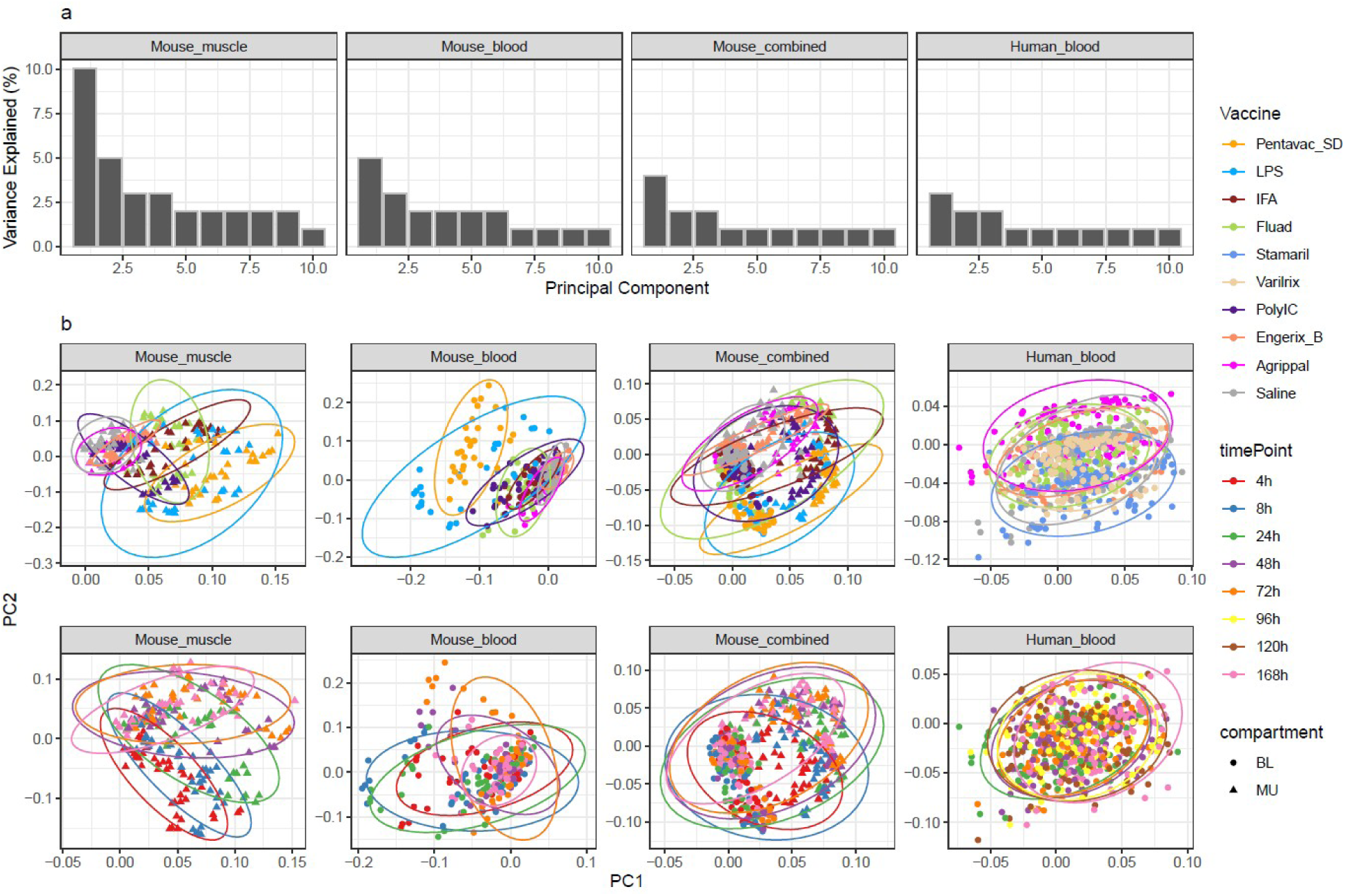
Principal component analysis of vaccine formulations across three transcriptomic datasets. **(a)** Percentage of variance explained by the principal components. **(b)** Sample projections onto the first two principal components, coloured by vaccine formulation (top) and by time point (bottom).

**Supplementary Figure 2:**
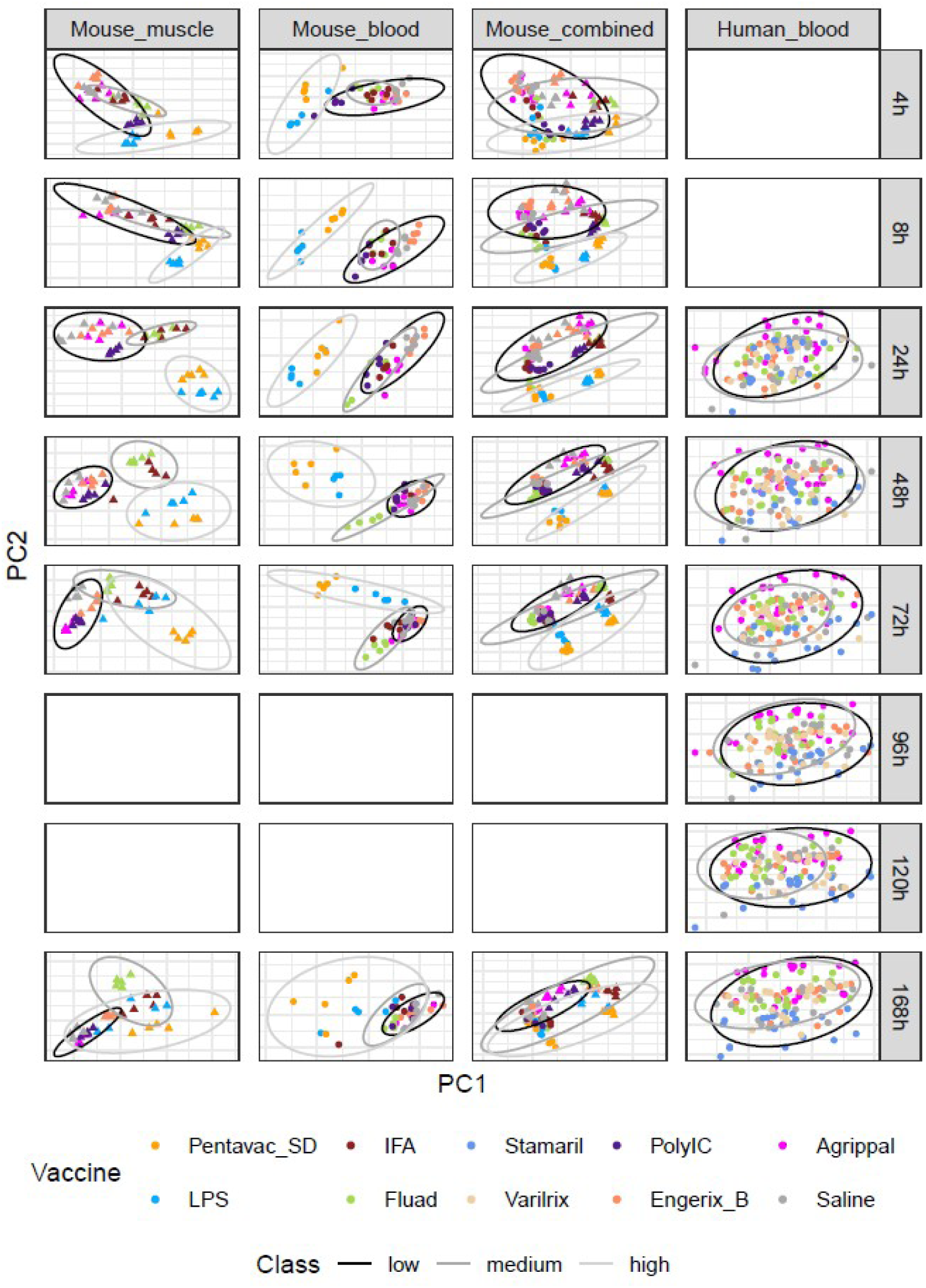
Principal component analysis of vaccine formulations by time point across transcriptomic datasets. Samples are projected onto the first two principal components, coloured by vaccine formulation, with ellipses representing reactogenicity classes.

**Supplementary Figure 3:**
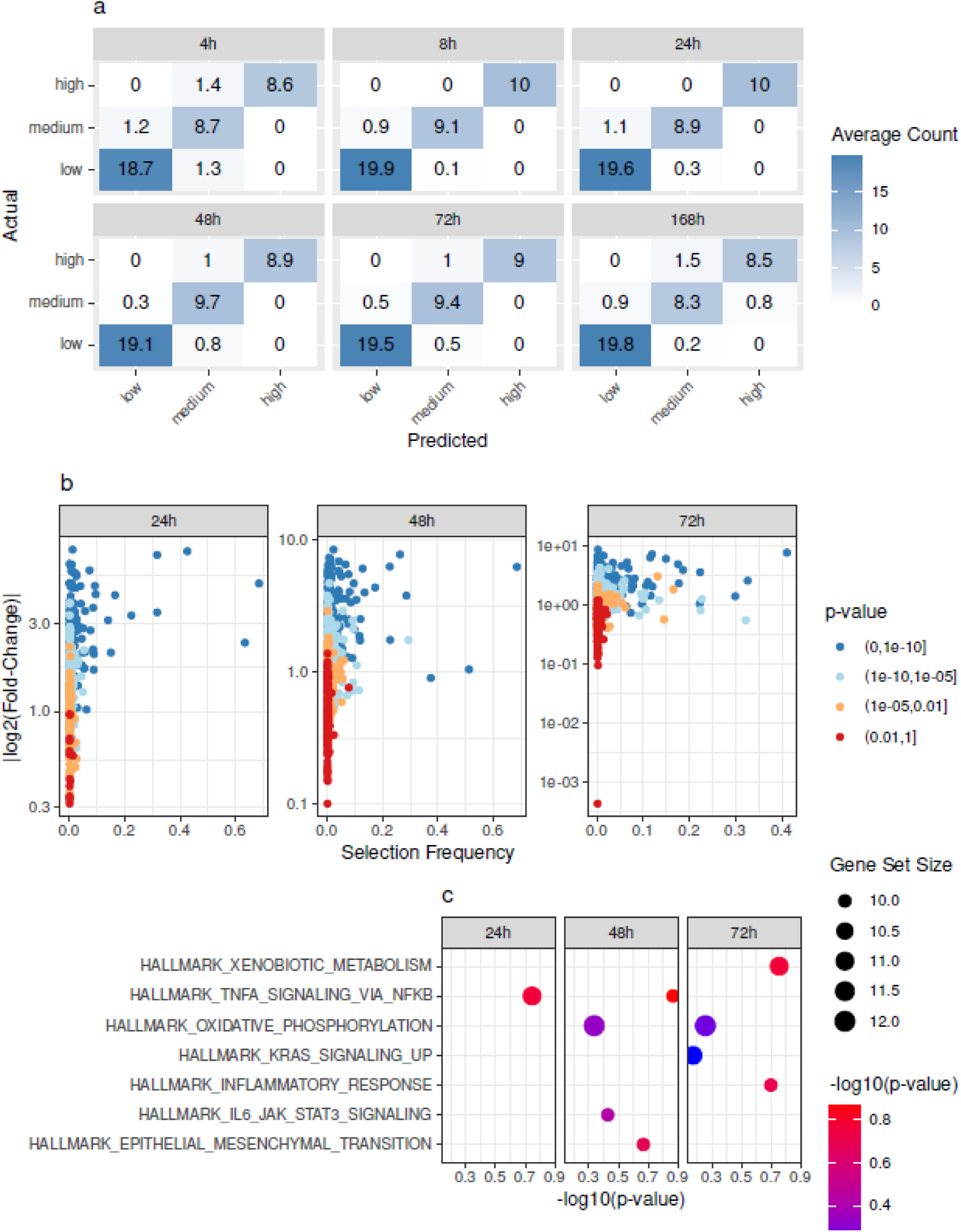
(a) Confusion matrices comparing predicted (columns) and observed (rows) reactogenicity classes in mouse muscle by time point. Results are averaged over 100 repetitions of nested cross-validation. **(b)** Comparison of gene predictive power (selection frequency computed via stability selection) and univariate differential expression (log fold change and p-value) between low and high reactogenicity classes at 24h, 48h and 72h. **(c)** Gene-set enrichment analysis on selection frequency, measuring the association between gene ranking and MSigDB gene sets.

**Supplementary Figure 4 :**
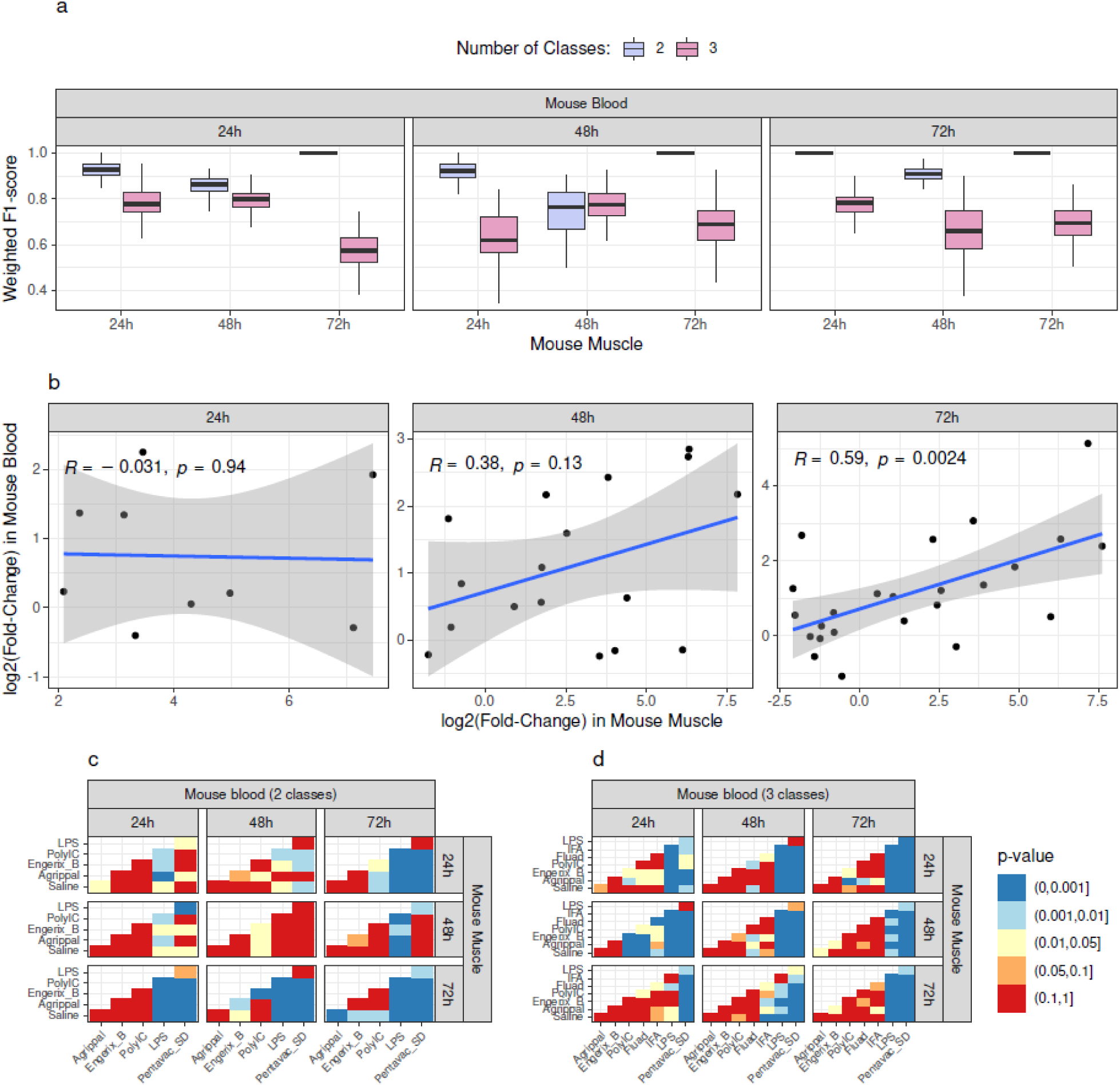
(a) Once trained on mouse muscle, the two and three-class models were evaluated on mouse blood at all combinations of time points using the weighted F1-score. **(b)** Scatter plots showing the relationship between log₂ fold-changes in mouse muscle and blood across low and high reactogenicity classes at 24h, 48h, and 72 h for the most predictive genes (selection frequency greater than 10%). **(c, d)** Adjusted *t*-test *p*-values for all pairwise comparisons of latent variables predicted at three time points (24h, 48h, and 72 h) using (c) two and (d) three-class models trained on mouse muscle and applied to mouse blood.

**Supplementary Figure 5 :**
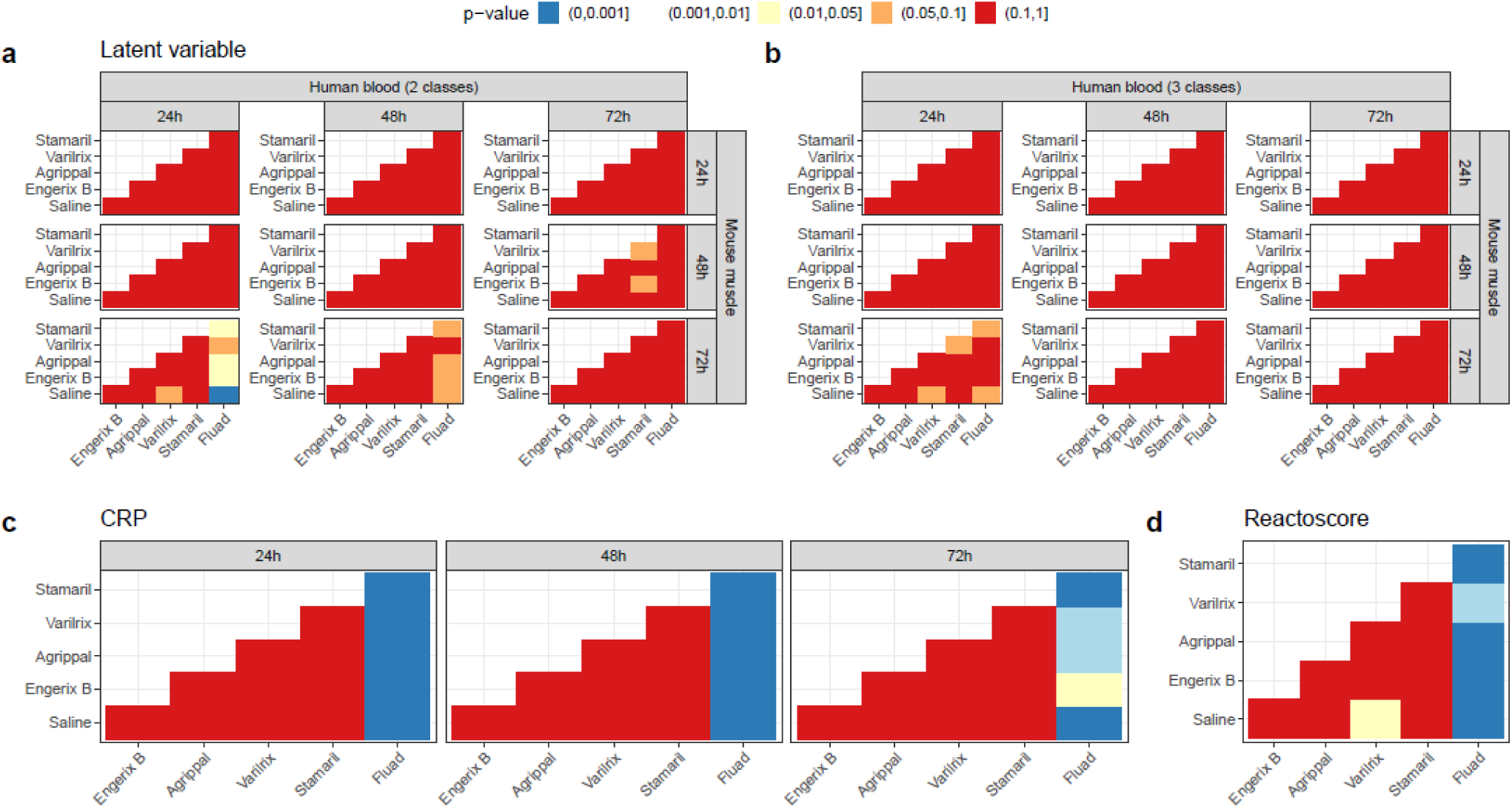
Adjusted *t*-test *p*-values for all pairwise comparisons of **(a)** latent variables predicted at three time points (24h, 48h, and 72 h) using two and three-class models trained on mouse muscle and applied to human blood; **(b)** CRP and **(c)** reactoScore values.

**Supplementary Table 1 :** Variable selection frequency computed with stability selection on mouse muscle at 24h, 48, and 72h.

**Supplementary Table 2 :** Differential analysis between low and high reactogenicity classes in mouse muscle.

**Supplementary Table 3 :** Adverse event classification. The isSpecific column indicates whether each term was considered specific to vaccination and therefore retained in the ReactoScore or excluded.

