## Supplementary figures and images for "A translational transcriptomic signature of vaccine reactogenicity for the evaluation of novel formulations"

### Supplementary Figure 1

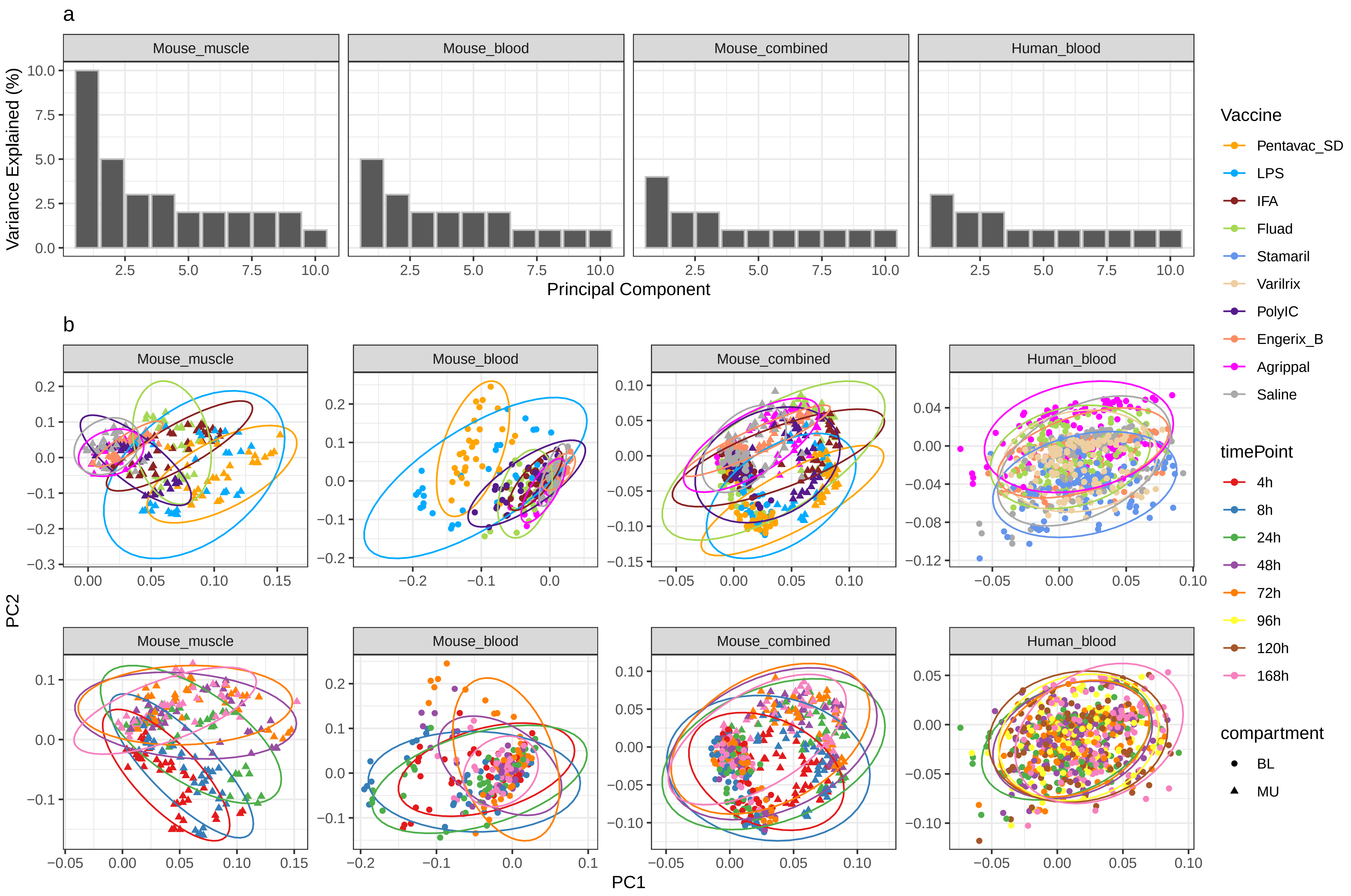

### Supplementary Figure 2

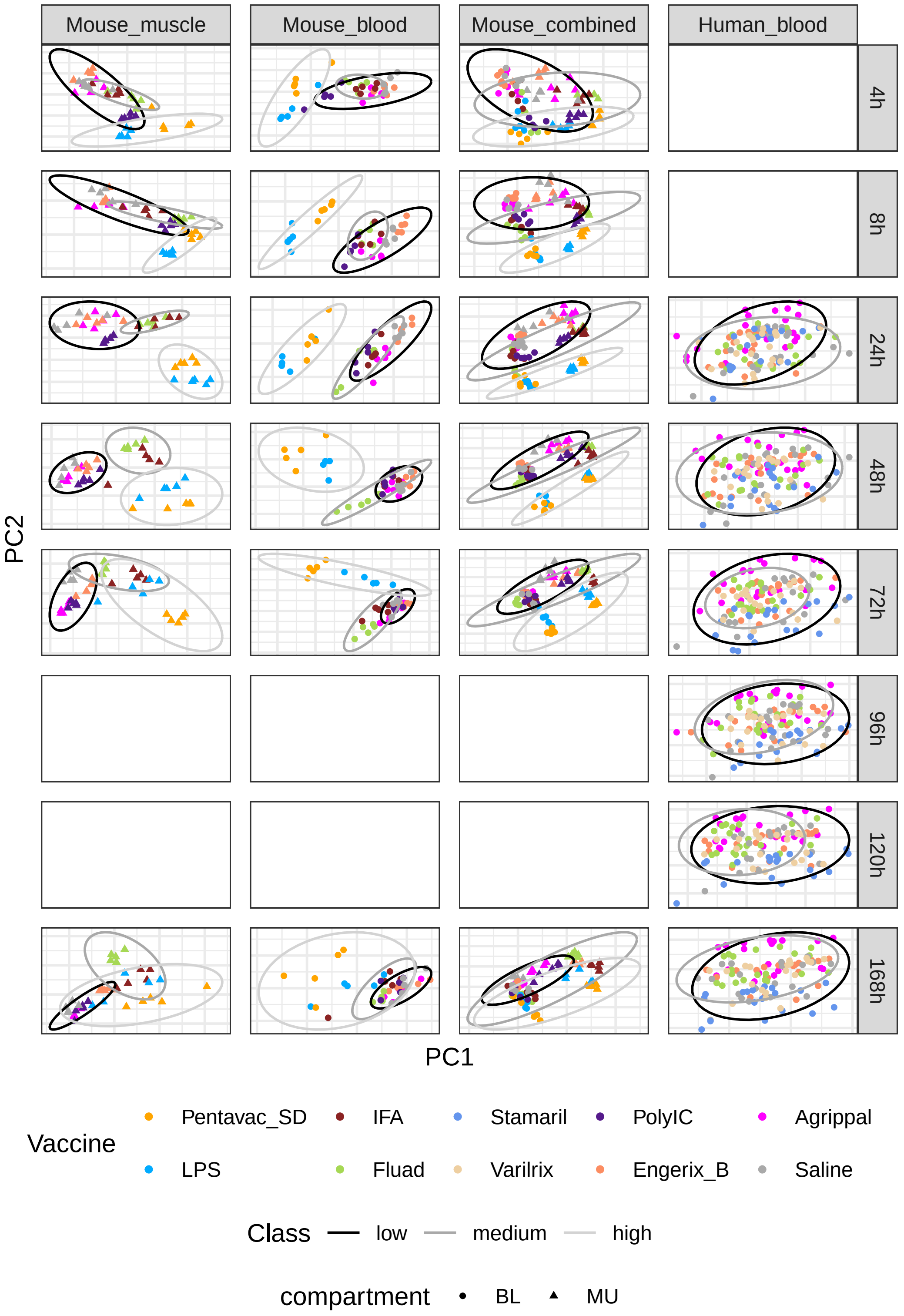

### Supplementary Figure 3

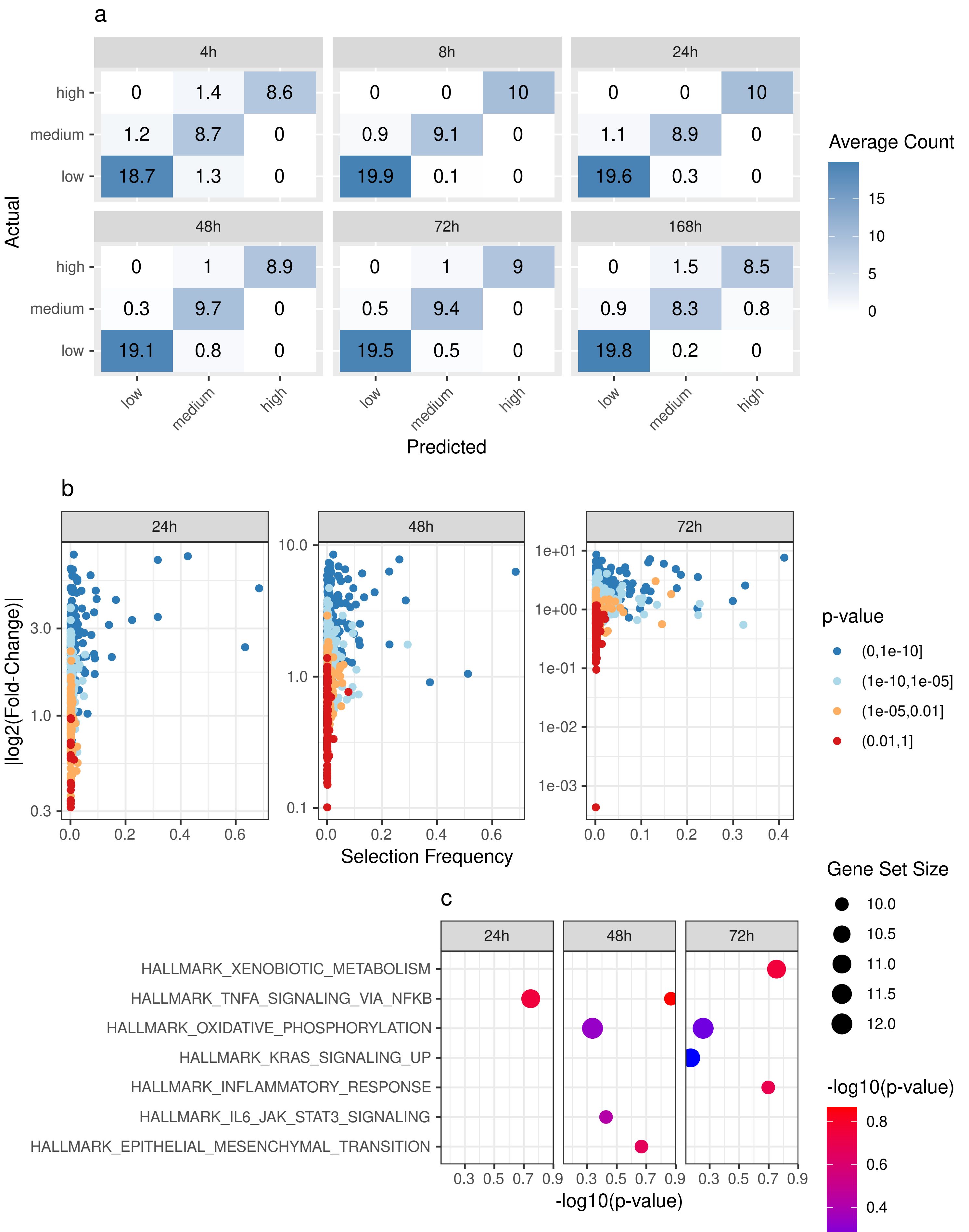

### Supplementary Figure 4

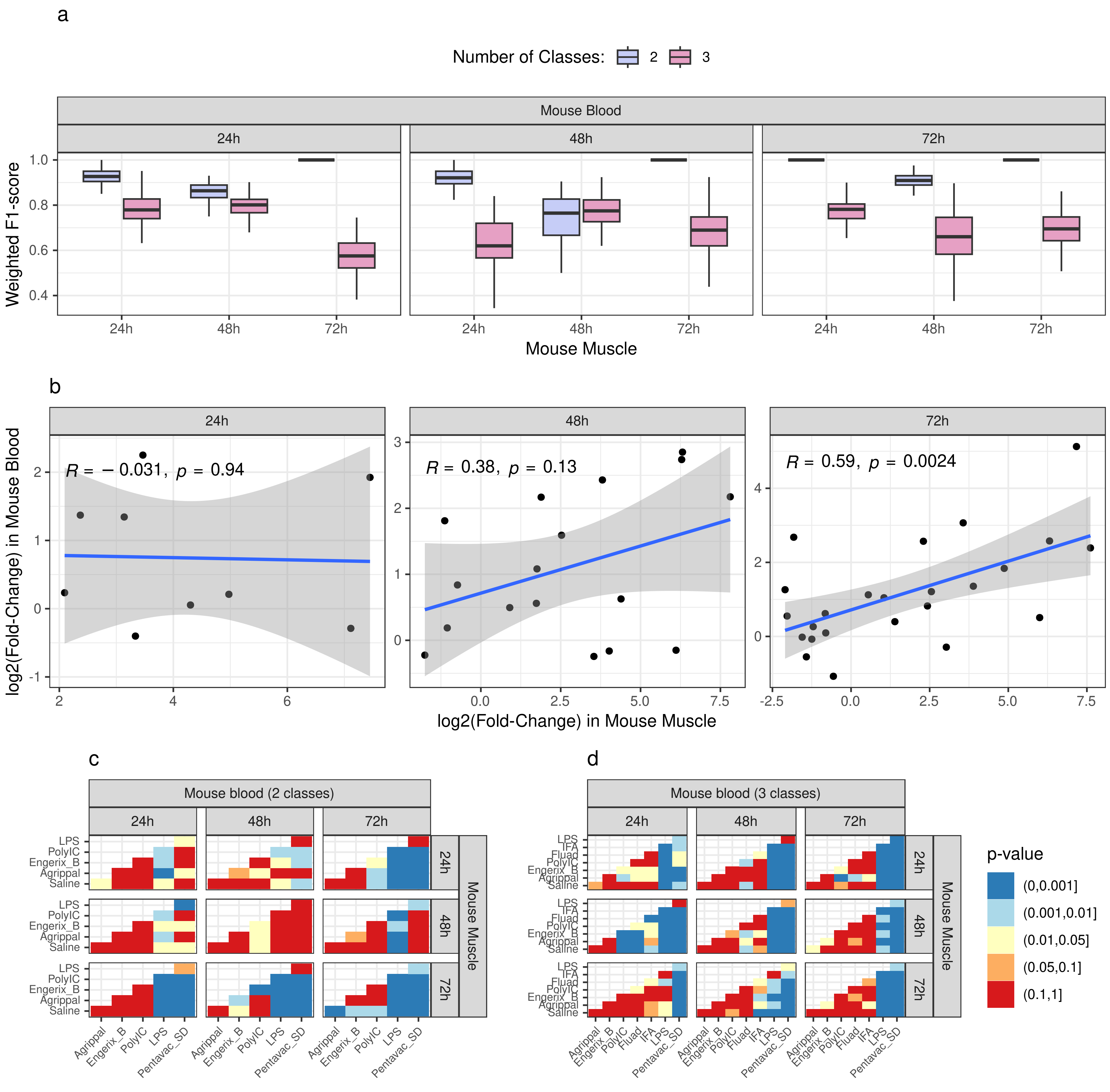

### Supplementary Figure 5

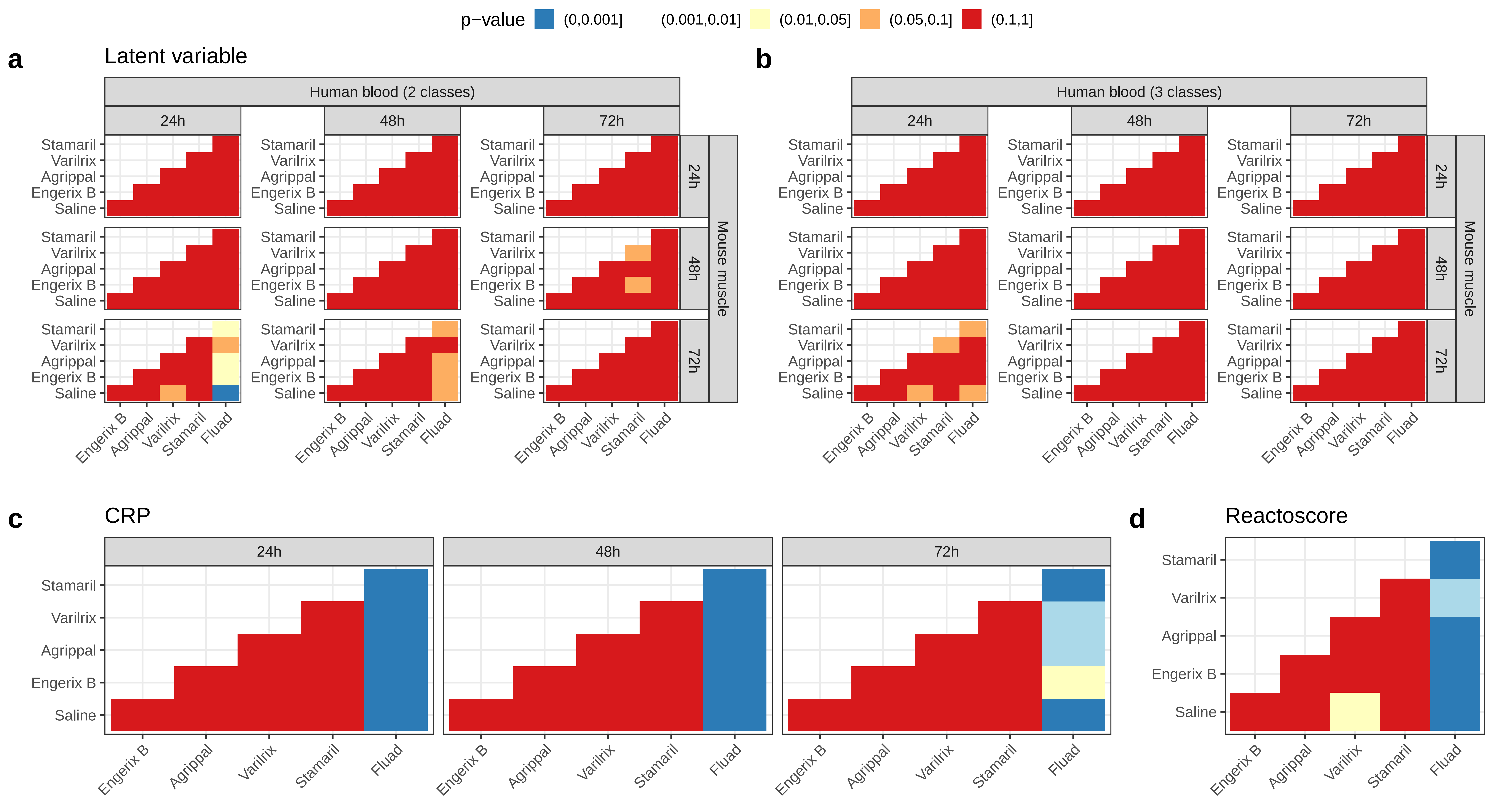
